# Visual environment of rearing sites affects larval response to perceived risk

**DOI:** 10.1101/2023.03.08.531703

**Authors:** Chloe A. Fouilloux, Jennifer L. Stynoski, Carola A. M. Yovanovich, Bibiana Rojas

**Affiliations:** University of Jyväskylä, Department of Biology and Environmental Science, P.O. Box 35, 40014, Jyväskylä, Finland; Clodomiro Picado Institute, University of Costa Rica, Coronado, San José, Costa Rica; School of Life Sciences, University of Sussex, Biology Road, BN1 9QG, Falmer, Brighton, United Kingdom; Department of Interdisciplinary Life Sciences, Konrad Lorenz Institute of Ethology, University of Veterinary Medicine Vienna, Savoyenstraße 1, 1160, Vienna, Austria

**Keywords:** larval vision, phytotelmata, poison frog, tadpole-rearing sites, predator-prey interactions, sensory ecology, limited visibility

## Abstract

1. Turbidity challenges the visual performance of aquatic animals. During development, environments with limited visibility may affect the fine-tuning of visual systems and thus the perception of, and response to, risk. While turbidity has frequently been used to characterise permanent aquatic habitats, it has been an overlooked feature of ephemeral ones.
2. Here, we use the natural diversity of ephemeral rearing sites (phytotelmata) in which the tadpoles of two poison frog species are deposited and confined until metamorphosis to explore the relationship between environments with limited visibility and response to perceived risk.
3. We sampled wild tadpoles of *Dendrobates tinctorius*, a rearing-site generalist with facultatively cannibalistic tadpoles, and *Oophaga* (formerly *Dendrobates*) *pumilio*, a small-phytotelm specialist dependent on maternal food-provisioning, to investigate how the visual environment in rearing sites influences tadpole behaviour. We hypothesised that turbid rearing conditions negatively impact both species’ ability to perceive risk, decreasing response strength to predatory visual stimuli. Using experimental arenas, we measured tadpole activity and space first on a black and white background, and then on either black or white backgrounds where tadpoles were exposed to visual stimuli of (potentially cannibalistic) conspecifics or potential predators.
4. When placed in a novel arena, the effects of rearing environment on *D. tinctorius* tadpoles were clear: tadpoles from darker pools were less active than tadpoles from brighter pools, and did not respond to either visual stimuli, whereas tadpoles from brighter pools swam more when paired with conspecifics versus odonate larvae, suggesting that tadpoles can visually discriminate between predators. For *O. pumilio*, tadpoles were more active on experimental backgrounds that more closely matched the luminosity of their rearing sites, but their responses to the two visual stimuli did not differ.
5. Larval specialisation associated with species-specific microhabitat use may underlie the observed responses to visual stimuli, which has implications for the stability of species interactions and trophic dynamics in pool communities. Together, our findings demonstrate that light availability of wild larval rearing conditions influences the perception of risk in novel contexts, and provide insight into how visually guided animals may respond to sudden environmental disturbances.

## Introduction

Sensory recognition is crucial for the success of predators and the survival of prey. While predators and prey are in an evolutionary arms race to refine their detection of each other, there are factors outside of their control that can limit the accuracy of their perception (*chemical contamination:* Weis and Candelmo 2012; *light pollution:* Minnaar et al. 2015). Nevertheless, developmental plasticity associated with the growth of organs and tissues creates the opportunity for bodily systems to optimise their function in a given environment. For example, the optic tecta of the brain in both fishes and amphibians enlarges when developing in habitats with higher conspecific densities (Gonda et al. 2009, Trokovic et al. 2011), while nutritionally poor diets in mammals (Lee and Houston 1993) and birds (Savory and Gentle 1976) induce an increase in gut size, optimising digestion. Sensory systems are particularly susceptible targets of selection, as they are energetically expensive structures (Niven and Laughlin 2008) whose performance is fundamental to individual survival and reproductive success (Streinzer et al. 2013). Bentho-pelagic elasmobranchs, for example, have denser and larger olfactory surfaces than benthic species, which supposedly compensates for the scarcity of visual cues in the open ocean (Schluessel et al. 2008). Male net-casting spiders (*Deinopis spinosa*) have enlarged principal eyes (and a more developed integration centre to process their inputs) compared to females and juveniles; the change in eye size in males is hypothesised to enhance predator detection as a result of switching from a sedentary to a wandering lifestyle (Stafstrom et al. 2017). These examples illustrate how sensory organs can compensate to adjust sensory function when faced with pressures from contrasting ecological niches (Schluessel et al. 2008).

Turbidity is a ubiquitous yet often underappreciated environmental pressure that shapes the dynamics of predator-prey interactions in aquatic environments (Abrahams and Kattenfeld 1997, Horppila et al. 2004, Van de Meutter et al. 2005). In experimental tests on the impact of turbidity, both background colour and the turbidity of rearing conditions have been found to influence the perception of risk by prey, affecting both their activity (invertebrates: Van de Meutter et al. 2005, fish: Leris et al. 2022) and space-use (invertebrates: Horppila et al. 2004), while generally reducing capture rates by predators (reviewed by Ortega et al. 2020). Laboratory studies have shown that the photic environment during development affects the fine-tuning of visual systems (*tadpoles:* Bridges 1970, *fishes:* Kröger et al. 1999, Fuller et al. 2010). From a proximate perspective, one mechanism that has been shown to flexibly change the spectral tuning of the eyes is the vitamin A1/A2 chromophore exchange system in the retina (Bridges 1972, Reuter et al. 1971). In this system, a lower A1/A2 ratio generates a red-shifted spectral sensitivity which can be advantageous in reddish environments (Corbo 2021) such as freshwater (Lythgoe 1979) and turbid waters (Jerlov 1976). Ultimately, while environmental conditions affect immediate (activational) individual responses, they also play a long-term (organisational) role in shaping the development of sensory systems throughout an individual’s lifetime (Snell-Rood 2013). Previous work on both killifish (Fuller et al. 2010) and guppies (Ehlman et al. 2015) has linked the developmental impact of limited light and turbidity with the spectral tuning of the visual system, which in turn has cascading effects on the behaviour of fishes in novel contexts. These studies establish an important connection between proximate-level restructuring of the visual system and ultimate-level changes in behaviour that can be used to predict outcomes of predator-prey interactions.

Phytotelm-breeding amphibians provide an exciting vertebrate model to further explore the relationship between environments with limited visibility and visually guided behaviours, as parents of some species transport their tadpoles to isolated pools of water in the vegetation where they are confined until metamorphosis. Phytotelmata are water-filled vegetal structures such as treeholes, palm bracts, and seed capsules, which are inhabited by diverse communities (Kitching 2000), and can appear as any combination of bright/dark and crystalline/turbid (Fouilloux et al. 2022a). Tadpole deposition sites are a unique system, because the conditions in lentic phytotelm environments can change suddenly and then persist for long periods of time (Leris et al. 2022, Fouilloux et al. 2021). In turn, the ‘darkness’ can be a result of light absorption by pigments and substances in the water itself, or the inner walls of the container, or both. Despite this remarkable variation, tadpole vision has never been explored in the context of the photic environment of rearing conditions. This is at least in part because earlier studies suggest that tadpole vision is generally ‘poor’ (e.g. myopic: Hoff et al. 1999, Mathis et al. 1988; low visual acuity expected on the basis of small eye size: Caves et al. 2018; Butler et al. 2022), and it had been well established that tadpoles rely heavily on chemical cues to navigate and assess their environment (Mathis and Vincent 2000, Weiss et al. 2021). More recently, however, experiments that isolate and combine sensory modalities have uncovered an important role of tadpole vision in conspecific communication and predator-prey interactions (Hettyey et al. 2012, Stynoski and Noble 2012, Kumpulainen 2022), and even some extent of illumination-dependent differential expression in vision related genes has been reported (Schott et al. 2022). Previous work demonstrating plasticity in tadpole vision substantiates the hypothesis that environments with selective light absorption may induce spectral shifts in their visual system (Donner and Yovanovich 2020, Schott et al. 2022) versus eliciting other adaptive responses, such as sensory compensation (Ehlman et al. 2015). Ultimately, although turbidity is frequently used to characterise permanent aquatic habitats, it has been an overlooked feature of ephemeral ones; as such, the impact of turbidity on the sensory development and visual recognition in species evolved to use ephemeral habitats remains unknown.

In this study, we investigated how wild-caught larvae of two phytotelm-breeding poison frog species (*Dendrobates tinctorius* and *Oophaga pumilio*), reared in pools with different optical characteristics, respond to predator and conspecific visual stimuli in novel contexts. *Dendrobates tinctorius* tadpoles are aggressive predators that will consume each other facultatively (Rojas 2014, Rojas and Pašukonis 2019, Fouilloux et al. 2022b). They are deposited in a wide range of microhabitat types and frequently cohabit pools with other tadpoles and insect larvae that can be either predators (e.g., dragonflies: Odonata) or prey (Rojas and Pašukonis 2019). In contrast, *O. pumilio* tadpoles depend on their mothers to deposit unfertilized, nutritive eggs in their pools (Stynoski et al. 2009), which serve as their primary food source throughout development (Brust 1993). In addition to being oophagous, *O. pumilio* tadpoles are small-pool specialists, primarily using bromeliad and other plant axils as pools, which they occupy singly; if multiple depositions occur, smaller tadpoles are cannibalised within a couple of days (Brust 1993, Stynoski 2009).

Given the variability in light availability and visibility inside phytotelmata, we hypothesise that the visual environment of rearing sites influences the development of sensory systems, and thus, the behaviour of larvae in novel conditions. Predatory tadpoles such as *D. tinctorius* may rely on vision (Kumpalainen 2022) and thus, low visibility during development could impact the phenotype of *D. tinctorius* visual systems (e.g., by red-shifting spectral sensitivity, Fouilloux et al. 2022a), which may mitigate the loss of acuity and improve predator performance in turbid conditions. With this said, a red-shifted spectrum in tadpoles is not a failsafe to overcoming the challenges of a visually limited environment. We predict that *D. tinctorius* reared in habitats with poor visibility would be generally less active (reduced prey response: Kimbell and Morrell 2015) and less discerning between visual stimuli (poor learning in turbid conditions: Chivers et al. 2012) than tadpoles reared in clear habitats. In comparison, given the restricted surface area of pools and high predation rates (67%, Maple 2002) in *O. pumilio* rearing sites, tadpole responses to perceived risk are better contextualised in terms of “seeking refuge and freezing” instead of the “escaping or attacking” response that more accurately describes *D. tinctorius’* behavioural repertoire. Thus, although *O. pumilio* tadpoles have been shown to respond differently to positive (food-bearing mother) and negative (predator) visual stimuli (Stynoski and Noble 2012), we expect a more generalised fear response where we predict that experimental contexts that are perceived as safer (i.e., more similar to rearing conditions) will elicit stronger behavioural responses (i.e., more activity) from *O. pumilio* tadpoles.

## Methods

### Study species

*Dendrobates tinctorius* and *Oophaga pumilio* are Neotropical poison frog species with intensive parental care. In *D. tinctorius* fathers care for the terrestrial clutches and transport tadpoles from oviposition sites to phytotelmata. *D. tinctorius* pool use is especially flexible compared to other phytotelm-breeding species, as these tadpoles occur in pools from a wide range of substrates that can be vastly different sizes (Fig. 1, volume: 19mL to 270L, height = 0- >20m, Fouilloux et al. 2021). Despite being facultative cannibals, *D. tinctorius* tadpoles are frequently placed in pools occupied by larger conspecifics (Rojas 2014). In these pools, tadpoles also encounter heterospecific tadpoles (generally *Allobates femoralis*) and insects (predatory Odonata larvae; Rojas 2014, Rojas and Pašukonis 2019, Fouilloux et al. 2021). For this study, *D. tinctorius* tadpoles were taken from dead palm bracts, treeholes, and fallen trees found around Camp Pararé, Les Nouragues Field Station, French Guiana from March to April 2022. Tadpoles were captured using small aquarium nets and spoons. Focal tadpoles ranged from stages 25 to 41 (Gosner 1960) and weighed 0.05–0.88 g.

**Figure 1.**
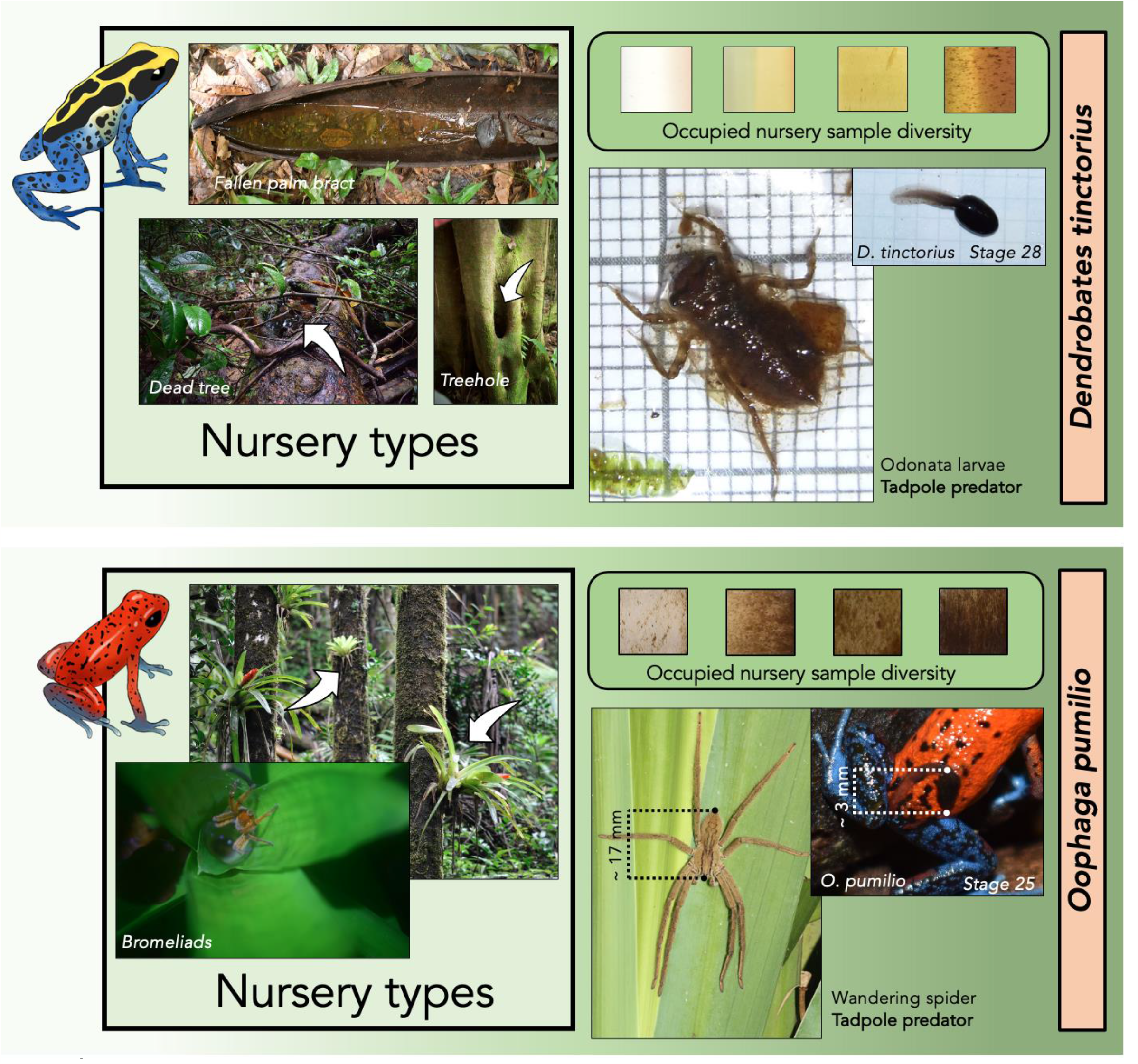
Characterization of the visual environment of two phytotelm-bound tadpole species. Schematic compares pool diversity in terms of both structure and turbidity, tadpole morphology, and frequently encountered predators. All photos by CF except for Odonata larvae (BR), forest epiphytes (US Forest Service - Southern Region, CC BY-SA 2.0) and spider (Stuart J. Longhorn, CC BY-SA 2.0). Line drawings kindly provided by Lia Schlippe Justicia.

Unlike in *D. tinctorius, O. pumilio* mothers are the primary caregivers, transporting tadpoles from terrestrial clutches to phytotelmata (Brust 1993). *O. pumilio* are much more specialised in their pool use than *D. tinctorius*, generally opting for small bromeliads and other axil-forming plants (volume: 10 to 100 mL, Stynoski *unpublished data*; Maple 2002). In addition to using much smaller pools, *O. pumilio* mothers feed deposited tadpoles with nutritive eggs, which serve as an obligate food source until metamorphosis (Brust 1993, Stynoski 2009). Tadpoles generally do not co-occur with other amphibian species. The predation rate of *O. pumilio* tadpoles is high, primarily as a result of snakes, aquatic beetle larvae (Family: Elateridae), and spiders (Family: Ctenidae, Maple 2002, Stynoski et al. 2014). Trials were conducted in La Selva Biological Station, Costa Rica from June to July 2022. *O. pumilio* tadpoles were captured from the field using pipettes and were tested the day of capture. Focal tadpoles’ stages ranged from 25 to 41 (Gosner 1960) and their weight from 0.01 to 0.12 g.

### Basic set-up

#### Background choice assay

All tadpoles (*D. tinctorius*, n = 20; *O. pumilio*, n = 25) were placed in the centre of an arena with a 2×2 black/white chequered background (Fig 2). The activity and background choice of tadpoles were quantified by scan-samples every 15 seconds for 15 minutes.

**Figure 2.**
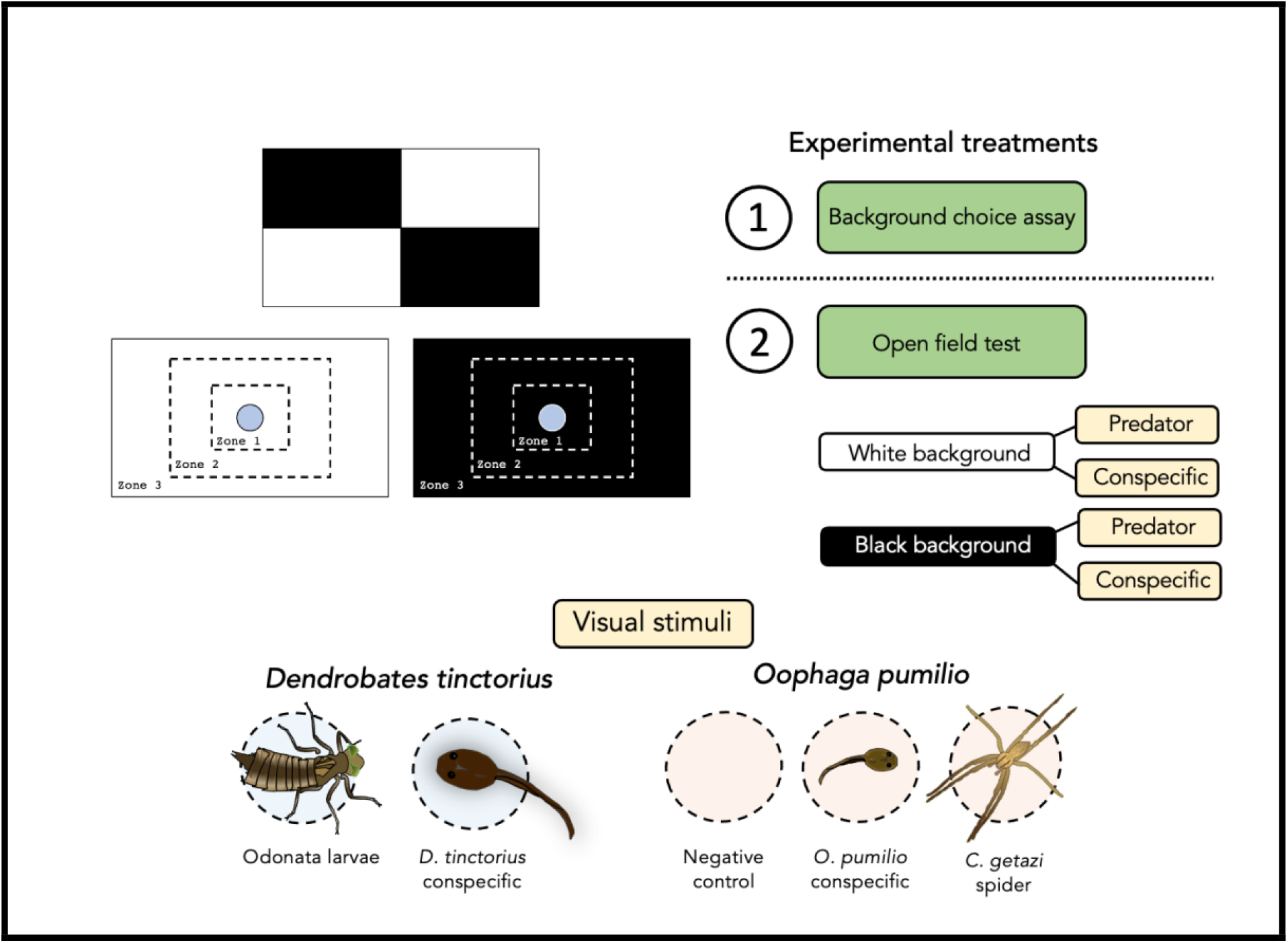
Schematic detailing experimental overview. The background choice assay was followed by open field tests with either white or black background treatments where tadpoles were exposed to either two or three visual stimuli in successive trials. Background order and the visual stimuli within each background block were randomised.

#### Open field tests

Following the background choice assay, we performed multiple open field tests. These consisted of an arena with either an entirely black or white background used as extreme cases of ‘crystalline dark’ and ‘crystalline bright’ visual environments, respectively (Fig. 1), containing different visual stimuli in the arena centre, i.e., either a conspecific tadpole or a known predator placed in an isolated container filled with rainwater. We recorded activity (swimming or not swimming) and space use (zone occupancy) of the focal tadpole at 15 second intervals for 15 minutes in each treatment. Zones were delineated by distances from the edge of the stimulus container in the arena centre, where Zone 1 was up to 2 cm away from the centre, Zone 2 was 2-5 cm away from the centre, and Zone 3 was beyond 5 cm.

We used a complete factorial design where all tadpoles were tested with all visual stimuli under both background conditions (Fig 2). Between individual stimulus trials within each background treatment, tadpoles were given five minutes to rest in a plastic container filled with rainwater. When changing between background colours, tadpoles were allotted at least 15 minutes of rest. After each 15 minute treatment, the water in the arena was changed to avoid the possibility of chemical-borne cues affecting tadpole behaviour.

To capture the variation of tadpole interactions in natural conditions, and also as a result of the limitation of available tadpoles, the size of the focal tadpole was at times smaller or larger than the tadpole that served as the visual stimulus. Mass differences between the focal and stimulus tadpoles ranged from −0.35g (where the focal tadpole was smaller than the stimulus) to 0.75g (where the focal tadpole was larger) for *D. tinctorius*, and from −0.10g to 0.054g for *O. pumilio*. Statistically, using the absolute mass of the focal tadpole or the mass difference between the focal and stimulus tadpoles predicted identical tadpole behaviour. All else being equal, we used focal tadpole mass instead of mass difference in all analyses for ease of interpretation (Supp Table 1, 2). For *D. tinctorius* experiments, in 18 out of 20 trials Odonata were the same size (n = 1) or larger (n = 17) than the focal tadpole (Supp. Fig 1).

Once all trials were finished, the experimental tadpole was weighed and staged. After each day of testing, all *D. tinctorius* tadpoles and odonate larvae were returned to their original pools. *O. pumilio* tadpoles were kept in the lab for an ongoing study. *Cupiennius getazi* or *C. coccineus* spiders were released at their capture point. All trials with each individual were completed on the same day. No tadpoles were used as a visual stimulus before serving as a focal tadpole. For *D. tinctorius*, ecological data about phytotelmata including information about conspecific count and the presence of heterospecific predators were available as a result of parallel monitoring surveys (see supplements for details of testing conditions, S1).

#### Standardised water sample photography, image analysis, and validation

Water samples from *D. tinctorius* rearing sites (n = 9 pools) were agitated and transferred to 1.5 mL glass vials and then photographed indoors on a white background next to a Macbeth XRite ColorChecker. Water samples from *O. pumilio* rearing sites (n = 25 pools) were transferred to a spectrophotometer cuvette and photographed in a lightbox (Puluz Ring Led Portable Photo Studio) on a white background next to a Macbeth XRite ColorChecker. In all cases raw photographs were taken using a Nikon D5300 digital camera (settings: ISO 500, f/13, shutter speed 1/125) from a 24 cm vertical distance (see supplements for details of sample collection and storage, S2).

All available water in *O. pumilio* phytotelmata was taken for spectrophotometry/photography measurements. Spectrophotometric readings of *O. pumilio* phytotelm water were analysed at the Clodomiro Picado Institute of the University of Costa Rica (see supplements for details on spectrophotometry (S3) and full transmittance spectra of *O. pumilio* pools (Supp. Fig 2)). Water spectra were taken in addition to standardised water sample photographs to validate the colour analysis of photos that were evaluated using ImageJ software (Abramoff et al, 2004). Such spectrophotometric quantification was not available for the *D. tinctorius* phytotelm water samples. Both methods aim to characterise light availability in the rearing sites to allow, albeit loosely, sorting of samples by visibility *sensu* Lythgoe 1979. For both *D. tinctorius* and *O. pumilio*, images were calibrated using the greyscale (standard reflectance set at 7% and 97%) and converted into .mspecs using the MicaToolbox plugin in ImageJ (Troscianko and Stevens 2015). Images were measured with objective camera vision, normalised by reflectance, and analysed as a linear normalised reflectance stack. Phytotelm samples were selected as regions of interest and their percentage of reflectance (relative to the standards) in the RBG spectra was computed, which then generated the reflectance of each colour channel relative to the standards.

### Statistics

Reflectance values from RGB photos (for both *D. tinctorius* and *O. pumilio*) were assessed using a principal components analysis (see Supplement S4 and Supp. Fig 3 for methods to compare photography and spectrophotometer quantification of water appearance (S4)). For both *D. tinctorius* and *O. pumilio* models the first component (PC1) explained more than 95% of the variance and thus were used as the predictor to represent differences in phytotelm photic environment (Fig 3, Supp. Fig 4). *O. pumilio* models were coded with either mean maximum transmittance from spectrophotometer readings or PC1 from photos (see Supplement S3 and Supp. Table 3). All statistical models were parameterized as count events; activity and space-use responses were predicted by experimental background, visual stimulus, the principal component (PC1) of microhabitat RGB % reflectances from standardised photography, and tadpole mass. *D. tinctorius* additionally included the conspecific count and the presence/absence (1/0) of natural predators in the original phytotelmata. All statistical models included tadpole identity (Tad_ID) as a random effect. *D. tinctorius* models also included phytotelm (Pool_ID) as a random effect as multiple tadpoles were sampled from the same phytotelmata. The best fitting interactive parameterization was assessed via a backward stepwise algorithm on a full three-way interactive model (interactions between experimental background, PC1, and visual stimulus were considered), followed by analysing the significance of two-way interactions by comparing model formulations via ANOVAs and assessing model ranks through AICc model selection. All models for both *D. tinctorius* and *O. pumilio* were coded using a negative binomial family. Final models passed quality checks for residual patterns, over/underdispersion, and zero-inflation using the simulation-based package “DHARMa” (Hartig et al. 2021). All models and figures were coded in R (v.4.1.1, R Core Team, 2022). For ease of comparison with *D. tinctorius*, the PC1 data from photos are reported in the results.

**Figure 3.**
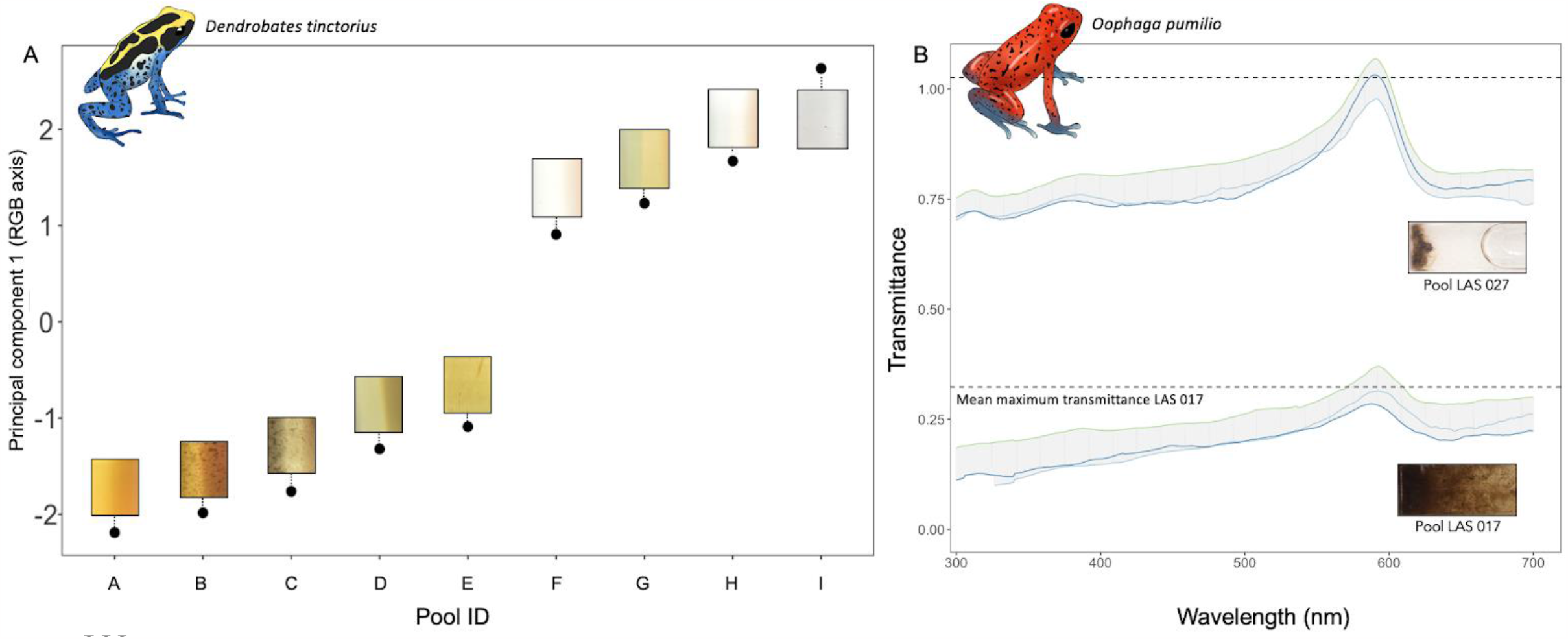
Diversity of tadpole pools with respect to photic environment. (A) The range of colour/brightness (PC1 with corresponding photographs, unedited) in pools occupied by *D. tinctorius*. (B) The variation in full absorbance spectra of *O. pumilio* microhabitats. Selected transmittance spectra show one of the darkest and one of the brightest pools (dashed lines: mean transmittance at the peak).

## Ethics statement

Experiments in French Guiana were approved by the DGTM (arrête No. R03.2022-01-07-00002), the Scientific Committee of Les Nouragues Ecological Research Station, and the Nouragues Nature Reserve (partnership agreement no. 01-2019 with BR). Experiments in Costa Rica were approved by the Resolución 292 of the Comisión Institucional de Biodiversidad de la Universidad de Costa Rica and SINAC-ACC-VS-16-2021 from MINAE, and ethics clearance CICUA-004-2021 from the Universidad de Costa Rica.

## Results

### Characterisation of phytotelmata and validation of photographs for the quantification of pool visual environment

Tadpoles of both *D. tinctorius* and *O. pumilio* occupy a range of phytotelmata that vary widely in the appearance of their water contents (Fig 1; Fig 3). They all have a red tint (see position of the transmittance peak in the red part of the spectrum, ∼600 nm in Fig 3 and Suppl Fig 2), such that the perceived differences lie on the bright-dark axis (see overall transmittance values in Fig 3 and Suppl Fig 2). For *D. tinctorius*, out of the nine pools sampled, three contained odonate predators and the total number of tadpoles ranged from one to ten. All 25 *O. pumilio* pools were singly occupied and none contained potential predators.

We validated the use of the reflectance values extracted from photographs of both species by comparing values with spectrophotometer readings from paired measurements of microhabitat samples from *O. pumilio* pools (see details in S4 of the supplements).

### Background choice assay

During the background choice assay, 75% of *D. tinctorius* tadpoles (n = 15) and 88% of *O. pumilio* tadpoles (n = 22) spent at least half of the allotted experimental time on black backgrounds. There was no apparent trend of an effect of tadpole mass or phytotelm photic environment on background choice for either species (Supp. Fig 5; Supp. Figure 6).

### Space use

Space use in *D. tinctorius* was predicted by both experimental background colour and visual stimuli, yet not in an interactive manner. Overall, tadpoles spent more time in the arena centre when on a white background (GLMM, z = 2.36 p = 0.018, Table 1) and when exposed to a conspecific versus an odonate larvae (GLMM, z = 3.07, p = 0.002, Figure 4A). Neither phytotelm photic environment, nor focal tadpole mass or predator presence in pools affected

**Table 1.** Model output for space use dynamics in *D. tinctorius*. Bolded values are statistically significant. The best fitting model was based on an additive parameterization.

| *D. tinctorius* space use (zone 1 counts) |  |  |  |  |
| --- | --- | --- | --- | --- |
| *Predictors* | *Log-Mean* | *SE* | *z value* | *p* |
| (Intercept) | 2.55 | 0.43 | 5.93 | **<0.001** |
| Condition [White] | 0.60 | 0.25 | 2.36 | **0.018** |
| PC1 | 0.10 | 0.11 | 0.94 | 0.347 |
| Predator [Conspecific] | 0.77 | 0.25 | 3.07 | **0.002** |
| Wild consp. count | -0.21 | 0.08 | -2.59 | **0.010** |
| Wild predator presence (Y/N) | 0.79 | 0.54 | 1.46 | 0.143 |
| Mass | 0.01 | 0.84 | 0.01 | 0.994 |
| **Random Effects** |  |  |  |  |
| $\sigma^2$ | 1.07 | | | |
| $\tau_{00}$ Pool_ID | 0.00 | | | |
| $\tau_{00}$ TadID | 0.18 | | | |
| $N_{Pool\_ID}$ | 9 | | | |
| $N_{TadID}$ | 19 | | | |

**Figure 4.**
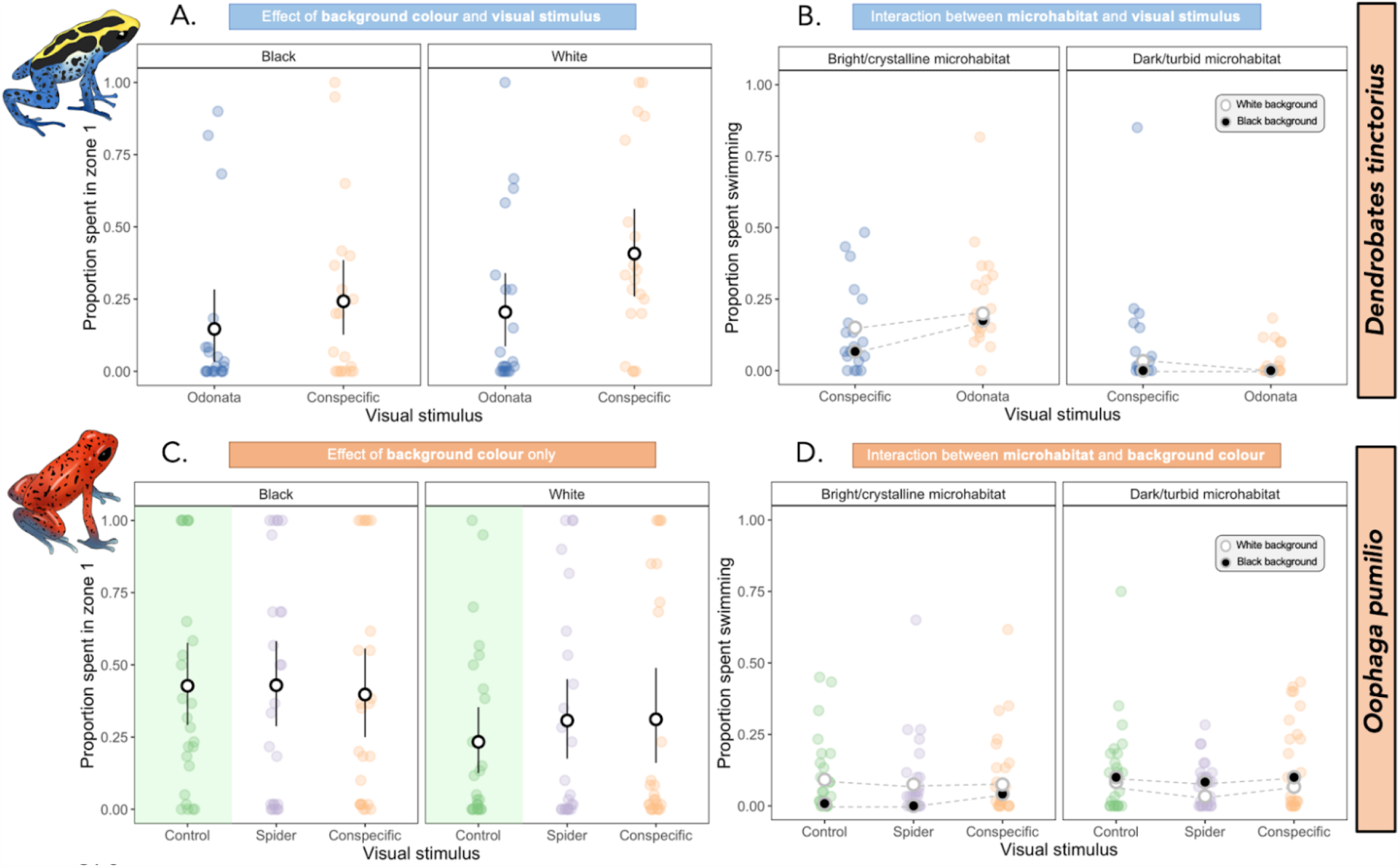
Space use and activity in poison frog larvae. The proportion of time spent in the arena centre is predicted by background colour and the visual stimuli in *D. tinctorius* (A) and only predicted by background colour for *O. pumilio* (C). Microhabitat influences activity in both species, where *D. tinctorius* is also influenced by visual stimuli (B) and *O. pumilio* tadpoles by background colour (D). Microhabitat photic environment is coded as a factor for visualisation purposes only, dashed lines help visualise slopes.

*D. tinctorius* space use (Table 1). Tadpoles from pools with more conspecifics spent significantly more time away from the arena centre (GLMM, z = −2.59, p = 0.010). For *O. pumilio*, only experimental background colour predicted space use: tadpoles spent significantly more time in the arena centre while on a white versus a black background (GLMM, z = −3.22, p = 0.001, Fig 4C). Neither visual stimuli, phytotelm photic environment or focal tadpole mass significantly influenced *O. pumilio* space use (Table 2).

**Table 2.** Model output for space use dynamics in *O. pumilio*. Bolded values are statistically significant. The best fitting model was based on an additive parameterization.

| *O. pumilio* space use (zone 1 counts) |  |  |  |  |
| --- | --- | --- | --- | --- |
| *Predictors* | *Log-Mean* | *SE* | *z value* | *p* |
| (Intercept) | 3.00 | 0.34 | 8.77 | **<0.001** |
| Condition [White] | -0.54 | 0.17 | -3.22 | **0.001** |
| Predator [Spider] | -0.03 | 0.20 | -0.17 | 0.862 |
| Predator [Cons.] | -0.10 | 0.20 | -0.48 | 0.629 |
| PC1 | 0.06 | 0.08 | 0.68 | 0.498 |
| Mass | 2.21 | 3.91 | 0.56 | 0.572 |
| **Random Effects** |  |  |  |  |
| $\sigma^2$ | 1.09 | | | |
| $\tau_{00}$ Tad_ID | 0.36 | | | |
| ICC | 0.25 |  |  |  |
| N Tad_ID | 25 |  |  |  |

### Activity

*D. tinctorius* tadpoles from darker pools swam less than tadpoles from lighter pools independently of the visual stimulus in the arena centre (GLMM, z = 3.71, p < 0.001, Table 3). We found a significant interaction between the visual stimulus and phytotelmata photic environment, where tadpoles from lighter pools (higher PC1 values) swam significantly more around conspecifics (GLMM, z = −2.19, p = 0.029, Fig 7) than around odonate larvae. We found no effect of experimental background colour, focal tadpole mass, number of conspecifics in microhabitats, or the presence of predators in microhabitats on *D. tinctorius* swimming behaviour (Table 3). *O. pumilio* tadpoles from lighter pools were significantly more active on white backgrounds than on black backgrounds (GLMM, z = 2.018, p = 0.044, Table 4, Fig 4D). These trends were consistent irrespective of the visual stimulus (Table 4). We also found that larger *O. pumilio* tadpoles were less active in general (GLMM, z = −3.622, p < 0.001, Supp. Fig 7b). A similar, yet non-significant, trend was found for *D. tinctorius* tadpoles (Supp. Fig 7a).

**Table 3.** Model output for activity in *D. tinctorius*. Bolded values are statistically significant. Best fitting model included an interaction between the visual stimuli and PC1 value representing microhabitat samples.

| *Predictors* | *D. tinctorius* activity (swim counts) |  |  |  |
| --- | --- | --- | --- | --- |
|  | *Log-Mean* | *SE* | *z value* | *p* |
| (Intercept) | 1.66 | 0.50 | 3.34 | **0.001** |
| Condition [White] | 0.34 | 0.21 | 1.59 | 0.111 |
| Predator [Conspecific] | -0.05 | 0.24 | -0.21 | 0.837 |
| PC1 | 0.53 | 0.14 | 3.71 | **<0.001** |
| Wild consp. count | 0.01 | 0.11 | 0.09 | 0.926 |
| Wild predator presence (Y/N) | 0.05 | 0.70 | 0.08 | 0.938 |
| Mass | -1.15 | 1.04 | -1.10 | 0.269 |
| Predator [Conspecific] * PC1 | -0.31 | 0.14 | -2.19 | **0.029** |
| **Random Effects** |  |  |  |  |
| $\sigma^2$ | 0.89 | | | |
| $\tau_{00}$ Pool_ID | 0.00 | | | |
| $\tau_{00}$ TadID | 0.37 | | | |
| $N_{\text{Pool\_ID}}$ | 9 | | | |
| $N_{\text{TadID}}$ | 19 | | | |

**Table 4.**
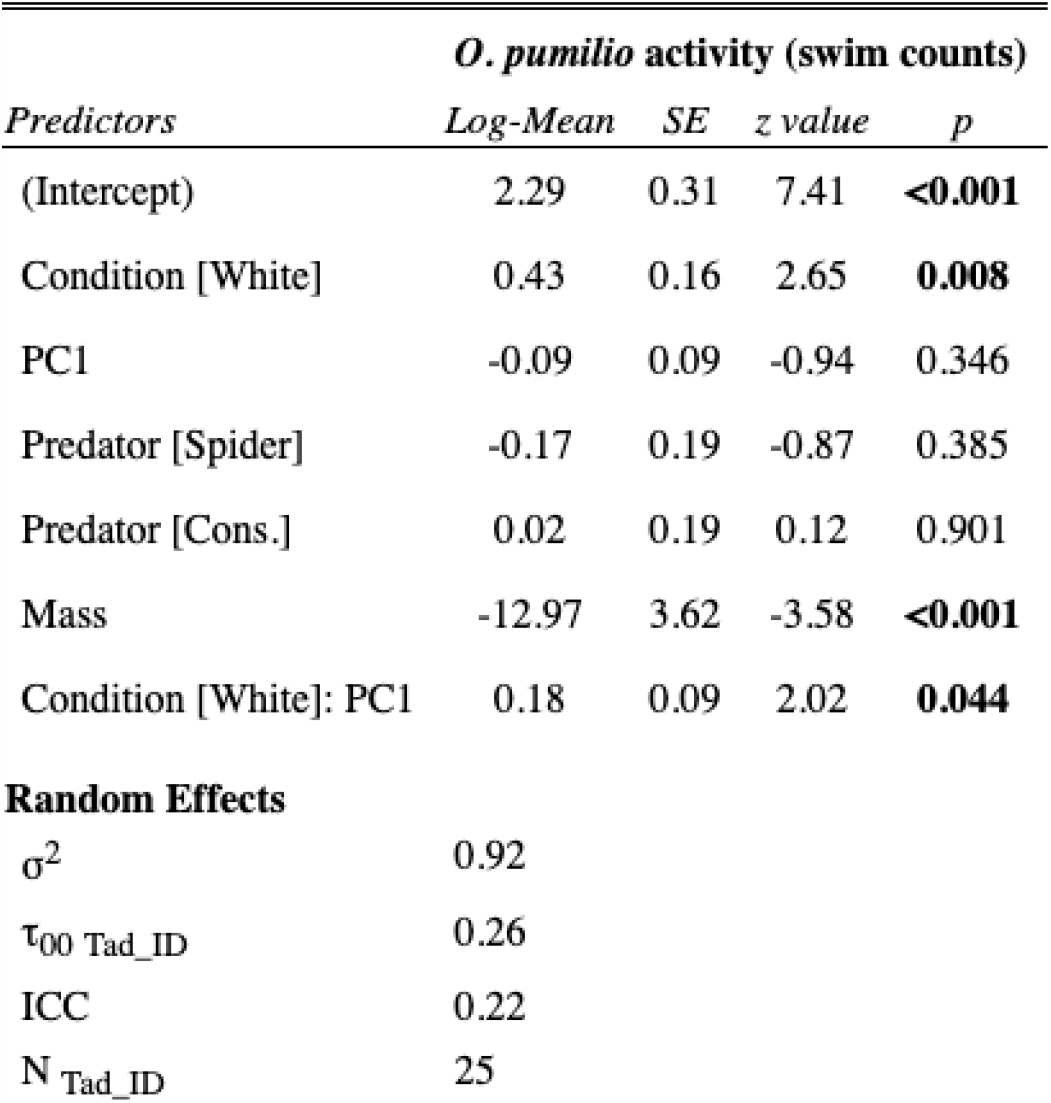
*O. pumilio* swimming activity model outputs. Bolded values are statistically significant. Best fitting model included an interaction between background colour and PC1 value representing microhabitat samples.

| *O. pumilio* activity (swim counts) |  |  |  |  |
| --- | --- | --- | --- | --- |
| *Predictors* | *Log-Mean* | *SE* | *z value* | *p* |
| (Intercept) | 2.29 | 0.31 | 7.41 | **<0.001** |
| Condition [White] | 0.43 | 0.16 | 2.65 | **0.008** |
| PC1 | -0.09 | 0.09 | -0.94 | 0.346 |
| Predator [Spider] | -0.17 | 0.19 | -0.87 | 0.385 |
| Predator [Cons.] | 0.02 | 0.19 | 0.12 | 0.901 |
| Mass | -12.97 | 3.62 | -3.58 | **<0.001** |
| Condition [White]: PC1 | 0.18 | 0.09 | 2.02 | **0.044** |
| **Random Effects** |  |  |  |  |
| $\sigma^2$ | 0.92 | | | |
| $\tau_{00}$ Tad_ID | 0.26 | | | |
| ICC | 0.22 |  |  |  |
| N<sub>Tad_ID</sub> | 25 |  |  |  |

## Discussion

Developmental conditions shape the way in which many organisms perceive risk. Throughout our sampling of phytotelmata we found that both *D. tinctorius* and *O. pumilio* tadpoles occur in a wide range of microhabitats that vary in their brightness and turbidity (Fig 1, 3) and that these differences have measurable effects on the tadpoles’ behaviour when paired with relevant visual stimuli.

### The effect of photic environment on activity across species

Both species of tadpoles use vision to assess their surroundings. Visual environment has different consequences on the behaviour of a tadpole predator versus one dependent on maternal provisioning throughout development. When assessing an individual’s preference for brightness, our choice experiments established a tadpole’s preference for black versus white backgrounds. Tadpoles of both species spent at least 75 percent of their time on black backgrounds where their dark brown bodies are considerably concealed. Similar results of animals choosing concealing environments have been shown in killifish (Kjernsmo and Merilaita 2012) and other amphibian larvae, all of which avoided non-concealing backgrounds (Eterovick et al. 2018) and had higher survival rates on concealing backgrounds when faced with predators (Espanha et al. 2015). Having established this preference, we followed up by introducing an additional risk component (predatory visual stimulus) on either entirely white or black backgrounds.

For both species, the visibility (including, but not limited to turbidity) of rearing conditions significantly influenced tadpole activity in a novel context. In *D. tinctorius*, tadpoles from darker pools moved less irrespective of the experimental background colour or visual stimuli (Fig 4, B) with which they were paired. This behavioural pattern has generally been established across the animal kingdom, where animals that were reared in more turbid conditions are less active and social in novel contexts (Fuller et al. 2010, but see work on damselfly responses reported by Van de Meutter et al. 2005). The implications of darker pools on activity are worth considering, as visually limited environments tend to negatively impact predation (Ortega et al. 2020). Predatory species such as *D. tinctorius* can consume both con- and heterospecifics (Rojas and Paškonis 2019); lower tadpole activity could have cascading effects on cannibalism rates, diet-quality, and thus time to metamorphosis (CF *unpublished data*, Kupferberg 1997) which may have negative implications for the survival of tadpoles, as pool stability is never guaranteed in ephemeral habitats (Fouilloux et al. 2021). From a community perspective, a shift in predator diet composition may affect the richness of the microhabitat, where species able to use other modalities to navigate their environment may outcompete vision-dependent species when faced with turbid conditions.

*O. pumilio* tadpoles were significantly more active on background conditions that more closely matched their rearing conditions (Fig 4, D). A similar trend was also found in Trinidadian guppies, where fish raised in clear water aquaria were more active in clear conditions compared to turbid conditions (Ehlman et al. 2015). Ultimately, the visual landscape of phytotelmata can drastically change depending on various biotic and abiotic factors, and it is not far-fetched to imagine a crystalline pool suddenly becoming turbid (or vice versa), as pools regularly dry and fill up again with clear rainwater and debris regularly falls from above (Romero et al. 2020). Based on our results, we would hypothesise that an *O. pumilio* tadpole from a brighter pool would consequently move less in dark conditions. This change in activity as a function of a changing environment has important implications for *O. pumilio* tadpoles, and may impact both the begging ritual between tadpoles and their mothers and the ability of tadpoles to detect and evade terrestrial predators entering their pools (Stynoski and Noble 2012, Stynoski et al. 2018). As most of *O. pumilio* tadpoles’ interactions occur at the air-water interface, manipulating object size, distance, and colour from the edge of a turbid versus a crystalline arena could disentangle the strength of tadpole responses to positive/negative stimuli and the acuity of tadpole vision in visually limited environments.

While we cannot conclude the behavioural changes in either species to be a result of visual restructuring, previous work has found lighting conditions of rearing environments (tea-stained vs. clear) to affect both foraging behaviour and opsin expression in the eye (Fuller et al. 2010). Ultimately, there exists a wealth of mechanistic oriented vision research, primarily in fish, showing that environmental light and colour affect the expression of visual pigments during development (Fuller et al. 2005, Shand et al. 2008). Thus, although we do not assess the visual system from a proximate perspective, multiple studies have shown that tadpoles rely on vision to assess risk (Hettyey et al. 2012, Szabo et al. 2021, this study), substantiating the role of vision as an important modality in the larval umwelt. While the link between turbidity as a particular case of ‘red-shifted’ environment (Jerlov 1976, Corbo 2021) and visual restructuring remains to be experimentally tested in tadpoles, recent interest in this field (Donner and Yovanovich 2020, Corbo 2021) is starting to yield exciting insights. It is well established that the challenge in turbid waters is not light availability (and thus it cannot be straightforwardly overcome with increased sensitivity) but rather distance-dependent contrast degradation due to non-image forming scattered light (Lythgoe 1979). In this scenario, myopic eyes such as those of tadpoles might not be so disadvantageous after all (Mitra et al 2022), as the image quality of the visual scene is proportionately better when viewing objects close by. There are neither theoretical nor empirical estimates of tadpoles’ visual acuity, however the recent confirmation that they tend to have spherical lenses even in species in which the adult lenses are flattened (Mitra et al 2022) opens up the possibility for tadpoles to present adaptations to maximise (within the constraints imposed by eye size) their spatial resolution and thus have a better chance to deal with turbid visual environments.

### Ecology shapes species’ responses to perceived risk

For each species, only one variable influenced consistently both activity and space use: the visual stimulus with which tadpoles were paired for *D. tinctorius*, and background colour for *O. pumilio*. The implication of these variables can be contextualised by comparing the biology of a habitat generalist versus a habitat specialist.

The outcome of most of *D. tinctorius* tadpoles*’* biological interactions is context-dependent, as *D. tinctorius* tadpoles can readily cannibalise conspecifics (Rojas 2014), but can also co-exist with them (Fouilloux et al. 2021); they can survive when odonate larvae are present, but can easily become their prey when the naiads are large enough (Rojas and Pašukonis 2019). Within the continuum of perceived risk for *D. tinctorius*, we find that in undesirable contexts (i.e., white background, an experimental environment tadpoles avoid when given the choice) a tadpole’s behavioural response is to “engage” with a potential threat. *D. tinctorius* tadpoles from clear pools swim more and spend more time in the arena centre where the stimulus is placed, suggesting that tadpoles recognize and seek to interact with conspecifics, even when provided only visual cues (similar to Kumpulainen 2022’s findings in a laboratory setting). In comparison, when paired with an odonate larva, which was larger or the same size as focal tadpoles 90 percent of the time, tadpoles from clear pools spend more time frozen at the edges of the arena. Previous work using open field tests associates elevated stress levels and risk aversion with arena centre avoidance (Champagne et al. 2010), suggesting that tadpoles might occupy the arena edges in response to the potential risk posed by the naiad. While *D. tinctorius* tadpoles from dark coloured pools seem to have adopted a “sit and freeze” strategy no matter the condition, tadpoles from lighter pools are able to recognise and change strategies depending on the visual stimulus with which they are faced. As a result of the flexibility of *D. tinctorius* deposition strategies, tadpoles are more likely to encounter diverse heterospecifics (tadpoles, dragonflies, mosquitoes) on varied backgrounds (dead bark, young plants, treeholes) and thus, need to be able to distinguish different types of visual stimuli.

These findings do not suggest that *D. tinctorius* tadpoles are more visually oriented than *O. pumilio* but instead highlight how the natural history of different species shapes their response to perceived risk. For example, the effect of previous predator exposure generating stronger responses in ensuing encounters has been shown before in tadpoles (via chemical cues, Fraker 2009). In *D. tinctorius*, focal tadpoles from pools containing more conspecifics spent less time in the arena centre (Supp. Fig 8), suggesting that the strength of stimuli avoidance may be informed by previous antagonistic experiences. In comparison, *O. pumilio* are obligately oophagous tadpoles which, in most instances, develop singly and in particularly small water volumes. In the context of their natural history, a tadpole’s response to risk is to seek refuge. In the wild, this can be done by diving down to the detritus at the bottom of a leaf axil, for example. Previous work has established that *O. pumilio* tadpoles see and respond to diverse visual stimuli (Stynoski and Noble, 2012, Stynoski et al. 2018); in the context of this study the potential effect of a visual stimulus may have been overridden by the design of the experiment itself, as tadpoles were in an arena larger than any of their natural deposition sites and with nowhere to hide (800mL in this study vs. ∼15 - 150mL in previous experiments; Stynoski et al. 2018, Khazan et al. 2019). While this design was intentional to create a comparable context to *D. tinctorius*, its role in contributing to a tadpole’s perceived risk should be acknowledged. Overall, *O. pumilio* tadpoles spend less time in the arena centre when on a white background compared to black backgrounds (even in the control instance when there is no stimulus, Fig 4, C). For an *O. pumilio* tadpole, it could be that when on white backgrounds the potential “refuge” provided by edge effects of the arena draws tadpoles away from the arena centre. In a novel context, tadpole “freezing” appeared to be the general strategy adopted by tadpoles; without a refuge to hide in there was no reason for individuals to modulate their activity in response to the distinct risks of each stimulus.

### Phytotelmata as models for environmental disturbance

Phytotelmata provide natural mesocosms to test and manipulate diverse habitat quality indicators on the development of animals. The natural diversity in phytotelm turbidity can be used as a small-scale model to predict how other aquatic animals may respond to sudden changes in habitat quality (Busse et al. 2018, Romero et al. 2020). Ecologically, predator or prey advantage in turbid conditions depends on the short-term adaptability of either player’s detection/evasion systems (Abrahams and Kattenfeld 1997); ultimately, here two visually guided species respond in ways that could negatively impact their survival in the form of decreased foraging or hindered access to parental care. Although the plasticity of the visual system may mitigate these effects to some degree, based on our data we would expect animals dependent on other modalities to better succeed in visually limited environments. To our knowledge, this is the first study to quantify phytotelm photic environment: we validate that digital photography (with a colour standard) generates the same ranking of pool brightness/turbidity as full spectral readings obtained with a spectrophotometer, demonstrating the accuracy of photography as a quantification tool for microhabitat visual environment and opening the door for future studies to pursue other turbidity-related questions independently of having access to expensive laboratory devices.

### Conclusions

We compare the perception of risk of a flexible predatory species to a microhabitat specialist with an adapted oophagous diet, testing both species’ responses to visual stimuli having developed in varied photic environments. The perception of risk in animals is context-dependent, and the light quality of rearing conditions affects the response to risk in novel contexts. As animals face increasingly disturbed habitats, these results highlight how animals with varied natural histories depend on vision and how they might respond to sudden environmental disturbances.

## Acknowledgements

We are grateful to the staff of CNRS Guyane (French Guiana) and La Selva (Costa Rica) for logistic support, and to the Nouragues research station (managed by CNRS), which benefits from “Investissement d’Avenir” grants managed by Agence Nationale de la Recherche (AnaEE France ANR-11-INBS-0001; Labex CEBA ANR-10-LABX-25-01). BR is a proud participant of a multi-researcher partnership with the Nouragues Nature Reserve (Convention Cadre de Partenariat N°01-2019) aimed at improving and spreading knowledge about amphibians. CAF has many people to thank for this work-- in FG thank you to Ria Sonnleitner and Lia Schlippe Justicia for their vital help in the quest of finding tadpoles in the most unexpected places! *Un très grand merci à* Patrick Chatelet, who provided much appreciated engineering assistance in support of the project. In CR, CAF would like to thank the “Jungals” Team without whom great science and mental welfare surely would have suffered. CY is very grateful to Kevin Doran for fruitful discussions about the intricacies of the RGB colour space. This study was funded by the Academy of Finland (No 345974 to BR), a ‘Mobility Grant’ from the University of Jyväskylä (to CF), the Max Planck Institute for Chemical Ecology/CONARE of Costa Rica (CVI-19-2021 to JLS), the International Centre for Genetic Engineering and Biotechnology (CRP/CRI19-04 to JLS), and European Union’s Horizon 2020 program (MSC-IF 101026409 to CY).

