## Supplemental Methods and Figures for "Visual environment of rearing sites affects larval response to perceived risk"

#### SUPPLEMENTARY METHODS

##### 1. Testing conditions

*D. tinctorius* trials (n = 20) were conducted outdoors under a white tarp to both prevent rainfall from affecting measures and to attempt to standardise ambient light above the arena (dimensions: 28.5 cm x 21.0 cm x 9.0 cm, 3mm thickness Rotho LOFT filled with 2.9 L of rainwater). *O. pumilio* trials (n = 25) were conducted in a laboratory with overhead light where temperature was held at 26°C (arena dimensions: 25 cm x 20 cm x 14.5 cm, 5 mm thickness plexiglass, filled with 800 mL of rainwater). After capture, tadpoles were housed singly and were given > 1 hour to acclimate before testing. All trials were conducted from 12-18h, when both of these diurnal species are naturally active.

##### 2. Collection of water samples from pools and storage of *O. pumilio* water samples

Water from occupied phytotelmata was collected and photographed on the same day that tadpoles were tested. *D. tinctorius* samples (n = 9 pools) were collected in syringes that were completely submerged in a disturbed pool. Samples, which ranged from 1.5 mL to a maximum of 10 mL, were then transported to the field station. For *O. pumilio,* after tadpoles were captured, nursery water was collected by submerging a pipette with the tip cut off into the bottom of the leaf axil and immediately transferred to a 1.5 mL glass vial (n = 25 pools). *O. pumilio* phytotelm water was photographed both the day of collection and the day of spectrophotometry measurements to confirm that samples had not changed or degraded between sampling and processing dates. After sampling and initial photography, water samples were refrigerated at 4°C until processing with a spectrophotometer at the Clodomiro Picado Institute of the University of Costa Rica.

##### 3. Spectrophotometry measurements of water from *O. pumilio* phytotelmata

For spectrophotometry measurements, *O. pumilio* phytotelm water samples were agitated and then diluted 1:5 with Milli-Q water before analysis, as pure samples led to absorbance levels above 1, at which point the samples are too concentrated and do not provide accurate absorbance outputs. We measured the full absorbance spectra of three technical replicates of each microhabitat sample, with the exception of three samples for which only two replicates could be measured due to insufficient volume. We used a Shimadzu (UV-1800) spectrophotometer with both ultraviolet range deuterium (D2, Type: L6380) and a 20W Tungsten halogen (W1, Type: NA55917) lamps with a silicon photodiode detector. The spectral absorbance of microhabitat samples was measured in quartz cuvettes (0.01m pathlength) against a Milli-Q water blank at 0.5 nm intervals. For ease of interpretation, absorbances were converted to transmittance values (Abs = log(1/T)). The maximum of the 300-700 nm full transmittance spectra were averaged between the three (or two) replicates for each pool (Fig 3A). This single value, the mean maximum transmittance, was then used as a representative value for phytotelm turbidity, where lower average transmittance represented darker/more turbid pools.

#### 4. Photography and spectrophotometry measures and validation

To verify the similarities in using photography and spectrophotometry to quantify optical characteristics of the water samples, spectrophotometer and photography values were compared by converting spectrophotometer spectra to RGB values based on human colour matching functions using the `spec2rgb` function from the R package "pavo" (Maia et al. 2019). Then, the relationship between the first principal components generated for spectrophotometer and photography RGB values (Supp. Fig 8) were compared by calculating Spearman's correlation coefficient *rho*. When comparing the correlation between the PC1 of the RGB values generated from spectrophotometer data and the RGB values of photographs of the same *O. pumilio* microhabitats, we found the two measures to be significantly and positively correlated (Spearman's *rho* = 0.84, p < 0.001, Supp. Fig 8). These data support the relationship between photography and spectrophotometry, and substantiate the accuracy of the reflectance values taken for *D. tinctorius* pools (Supp. Fig 8).

### SUPPLEMENTARY TABLES AND FIGURES

**Supplementary Table 1.** *D. tinctorius* activity or space use is not predicted by mass difference between tadpoles or focal tadpole mass. Model parameterizations yielded identical results (significant variables highlighted), were all within 2 AICc of each other, and passed all suitability checks using the DHARMa package (residuals, dispersion, inflation). All models included random effect structure described in main text.

Statistically significant predictors highlighted

| | Response | Final model structure | AICc | DHARMa Check |
|---|---|---|---|---|
| Absolute mass (Mass) | Zone 1 | Background + Predator + PC1+ #conspecifics + Predator (Y/N) + **Mass** | 509.06 | Pass |
| Mass Difference (Focal-Center) | Zone 1 | Background + Predator + PC1+ #conspecifics + Predator (Y/N) + **Mass difference** | 509.76 | Pass |
| Absolute mass (Mass) | Swim | Background + Predator * PC1 + #conspecifics + Predator (Y/N) + **Mass** | 440.36 | Pass |
| Mass Difference (Focal-Center) | Swim | Background + Predator * PC1 + #conspecifics + Predator (Y/N) + **Mass Difference** | 440.36 | Pass |

**Supplementary Table 2.** Model parameterizations with either mass difference between tadpoles or focal tadpole mass for *O. pumilio* yielded identical results (significant variables highlighted). Neither mass difference nor absolute mass was significant in predicting tadpole space use, but did play a role in overall activity. All models passed suitability checks using the DHARMa package (residuals, dispersion, inflation). Models included random effect structure described in main text.

Statistically significant predictors highlighted

| | Response | Final model structure | AICc | DHARMa Check |
|---|---|---|---|---|
| Absolute mass (Mass) | Zone 1 | Background + Predator + Mean Absorbance + **Mass** | 1140.03 | Pass |
| Mass Difference (Focal-Center) | Zone 1 | Background + Predator + Mean Absorbance + **Mass difference** | 1139.65 | Pass |
| Absolute mass (Mass) | Swim | Background * Mean Absorbance + Predator + **Mass** | 823.14 | Pass |
| Mass Difference (Focal-Center) | Swim | Background * Mean Absorbance + Predator + **Mass difference** | 818.35 | Pass |

**Supplementary Figure 1.** Effect of size difference between focal and Odonata stimulus in *D. tinctorius* tadpoles.

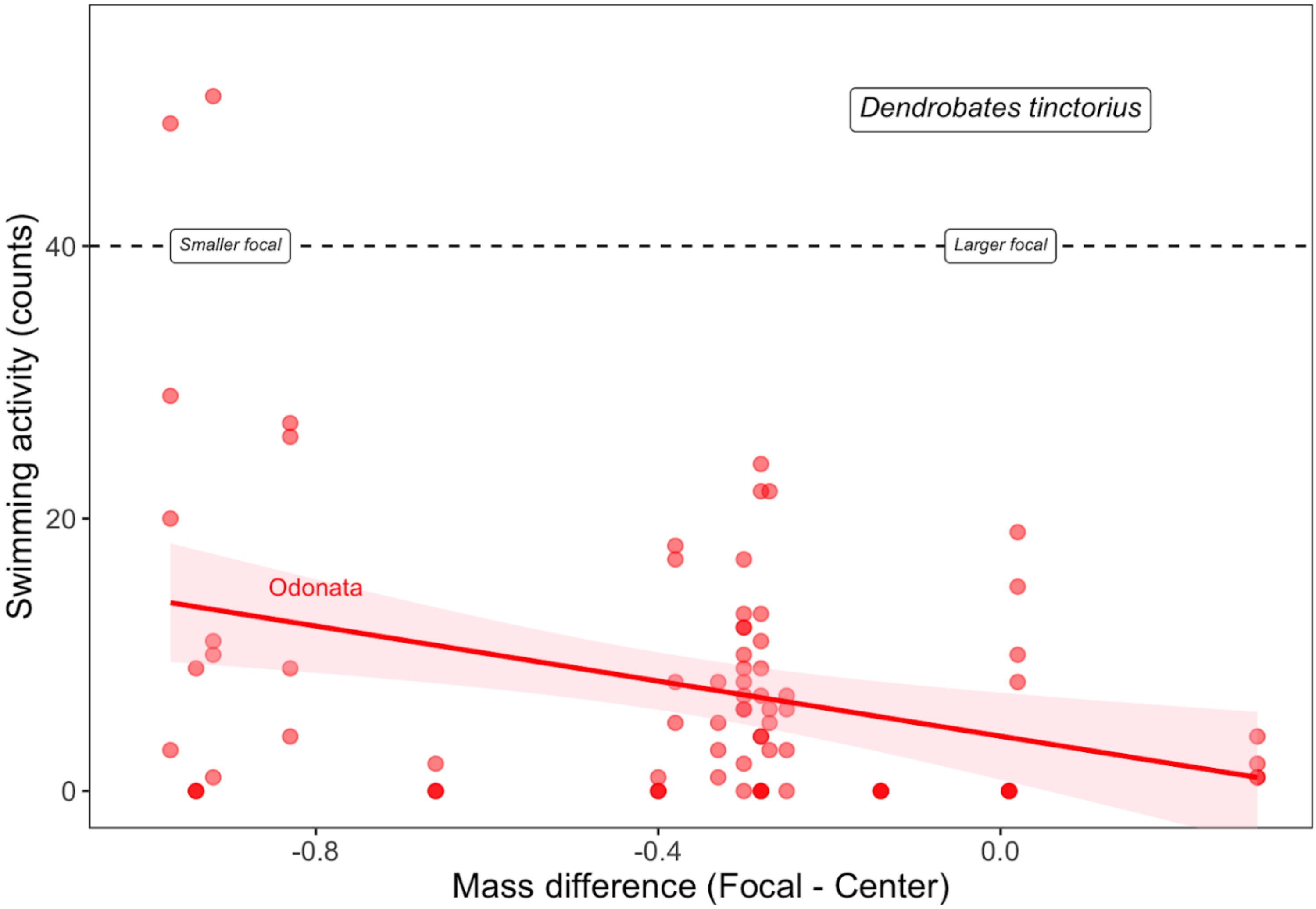

**Supplementary Figure 2.** The full transmittance spectra from spectrophotometer readings of each of the sampled microhabitats from La Selva, Costa Rica. Each panel represents an individual pool. Facets are ordered by increasing mean maximum transmittance.

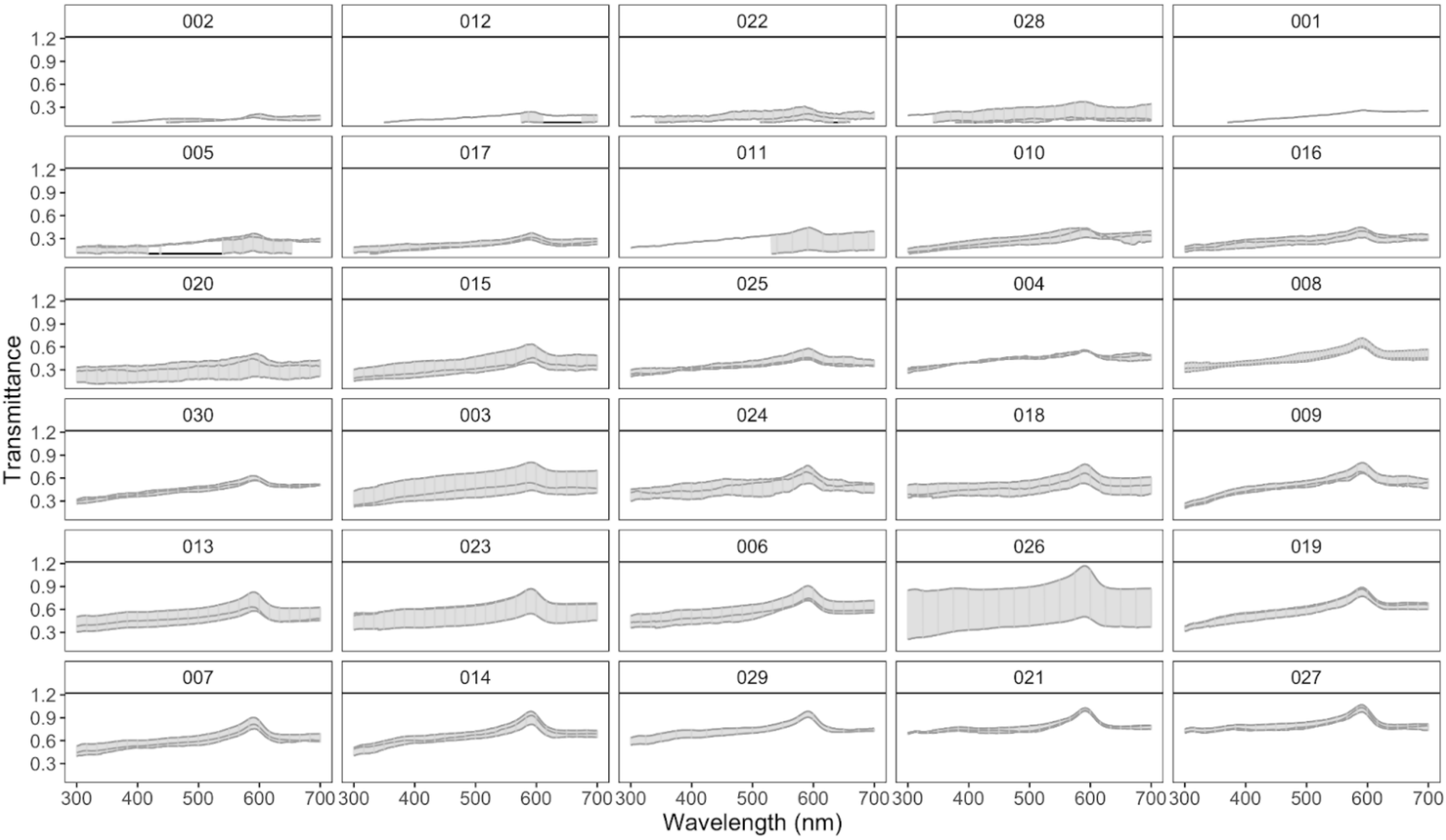

**Supp. Figure 3. Comparison between spectrophotometry and photography RGB values.** Panel (A) shows microhabitat transmittance spectra and their corresponding RGB values generated by the `spec2rgb` function from the "pavo" package. Panel (B) compares PC1 values generated from RGB values from both spectrophotometer and photography data. Second order polynomial fit to data with shaded regions representing the 95 CI using a "gam" smoothing function.

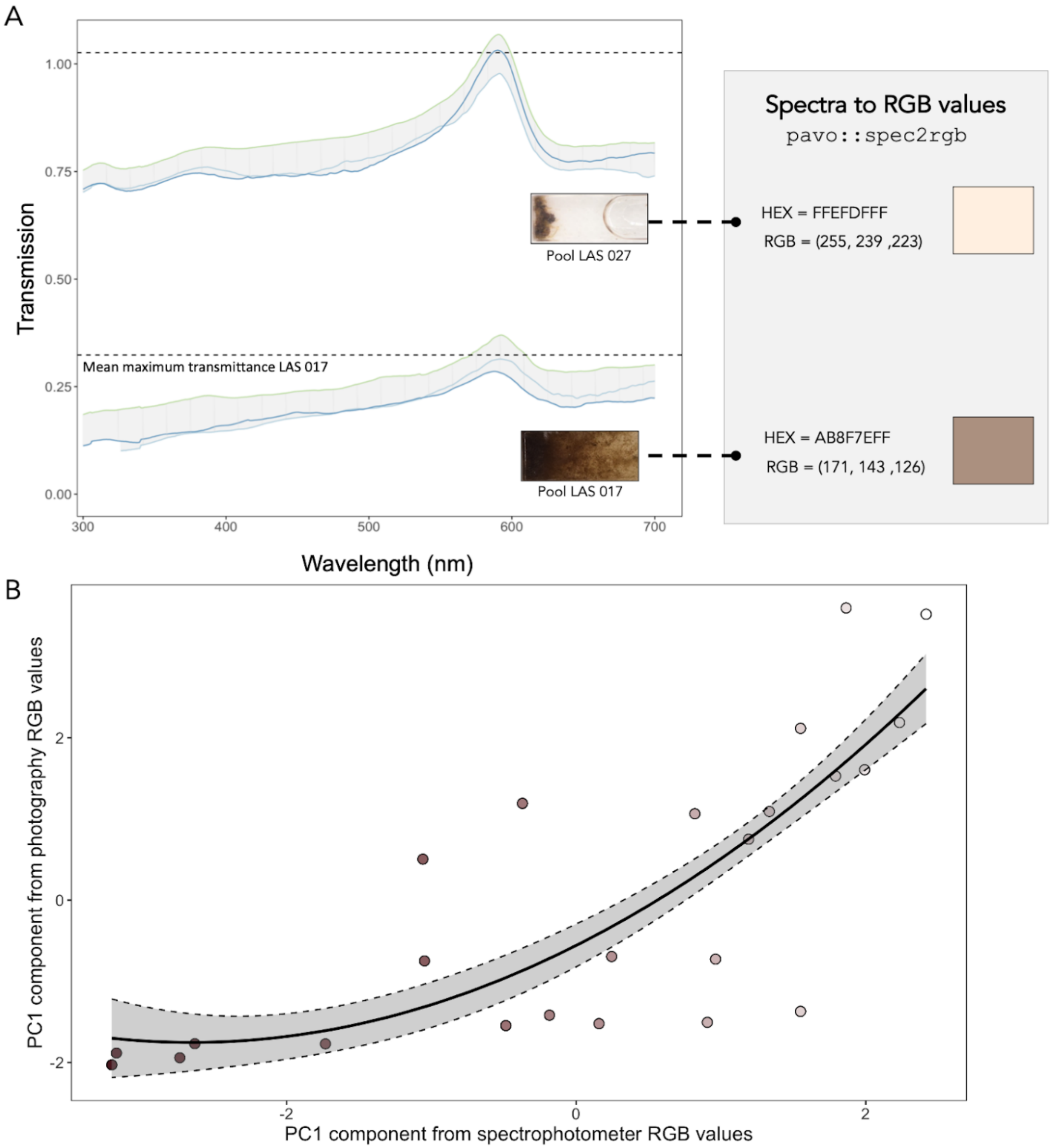

**Supplementary Table 3.** Model parameterizations with either spectrophotometry or photography data for *O. pumilio* yielded identical results (significant variables highlighted). All models passed suitability checks using the DHARMa package (residuals, dispersion, inflation). Models included random effect structure described in main text.

Statistically significant predictors highlighted

| | Response | Final model structure | AICc | DHARMa Check |
|---|---|---|---|---|
| Mean Max. Trans. | Zone 1 | Background + Predator + Mean Absorbance + Mass | 1140.03 | Pass |
| PC1 | Zone 1 | Background + Predator + %R-channel reflectance + Mass | 1142.08 | Pass |
| Mean Max. Trans. | Swim | Background * Mean Absorbance + Predator + Mass | 823.14 | Pass |
| PC1 | Swim | Background * %R-channel reflectance + Predator + Mass | 821.14 | Pass |

**Supplementary Figure 4.** Principal component 1 (PC1) explaining the RGB values from photos of each pool occupied by *Oophaga pumilio* tadpoles. We see that negative PC1 values capture more turbid pools while positive values show more clear pools. The same trend is found in *Dendrobates tinctorius* pools (main text Fig 3).

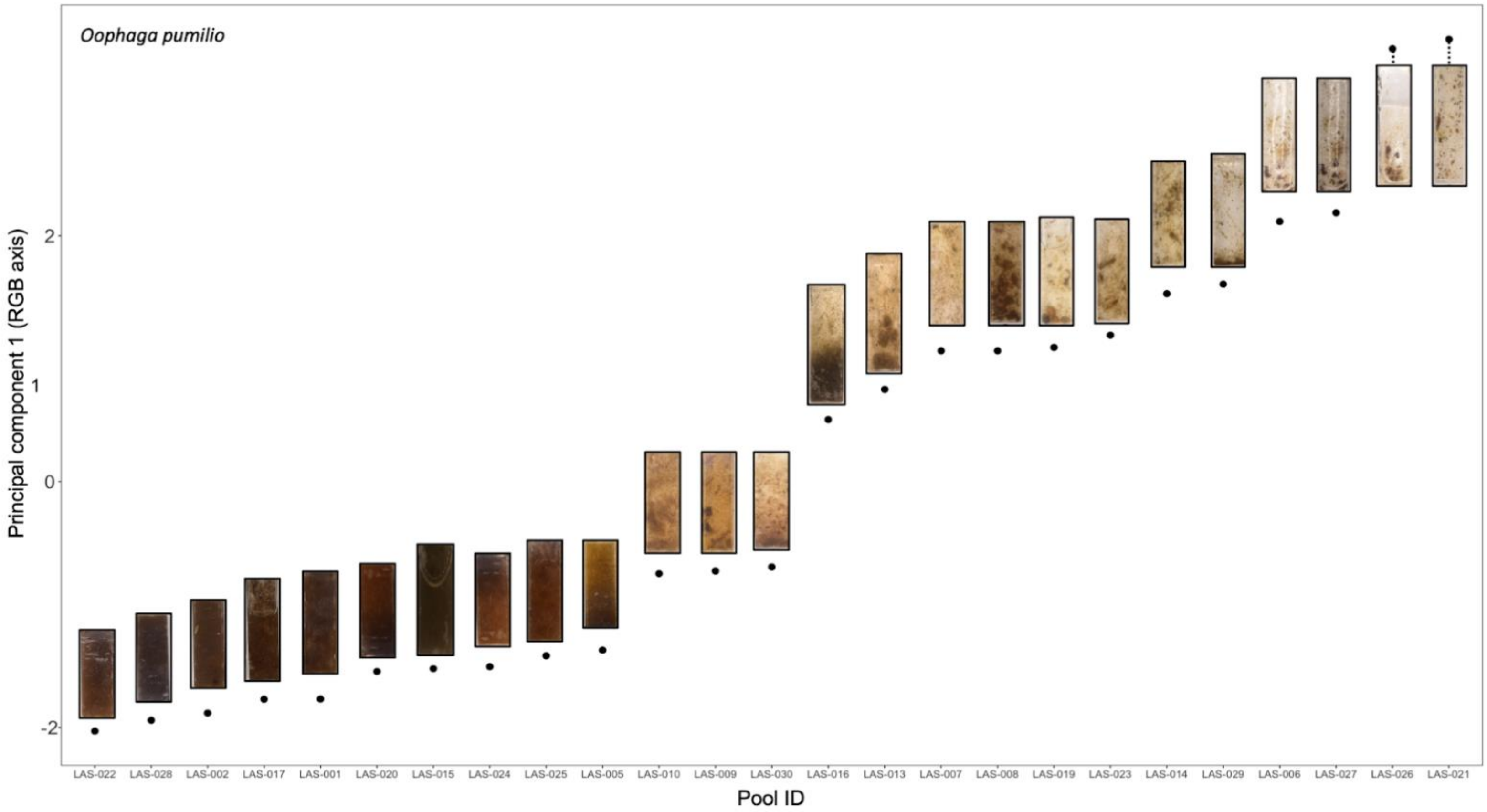

**Supplementary Figure 5.** Proportion of time spent on black background for *D. tinctorius* tadpoles. Points coloured by mean RGB PC1 from photos.

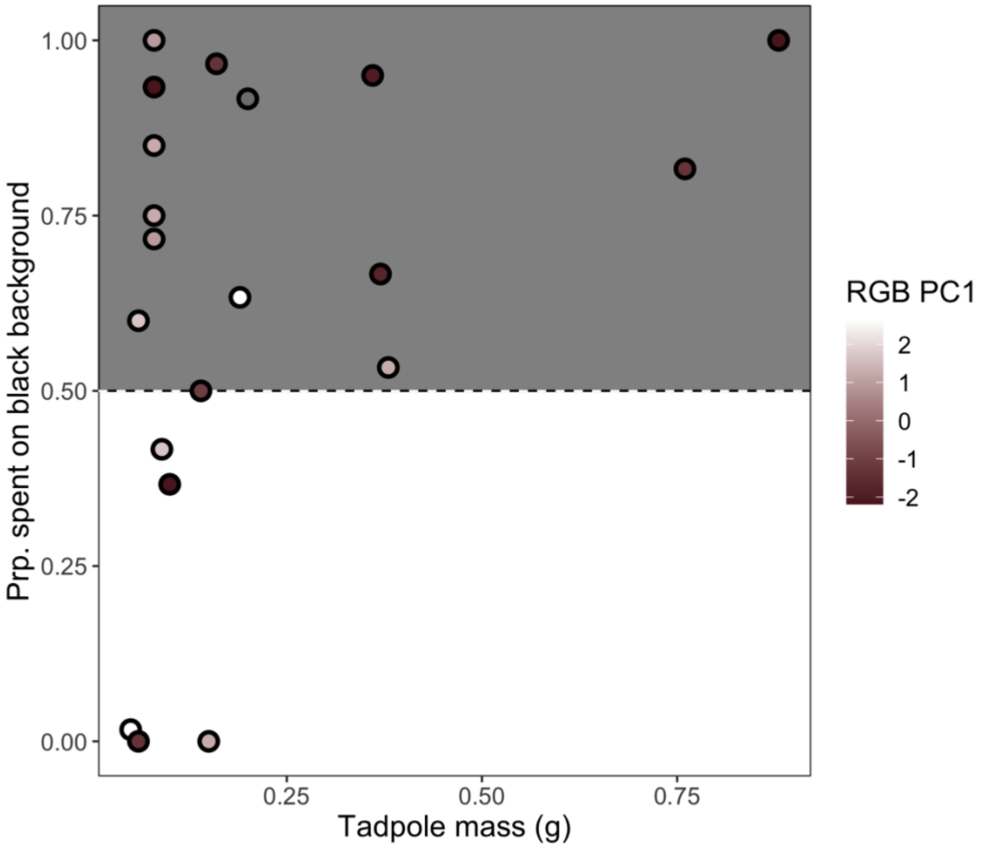


**Supplementary Figure 6**. Proportion of time spent on black background for *O. pumilio* tadpoles. Points coloured by mean RGB PC1 from photos.

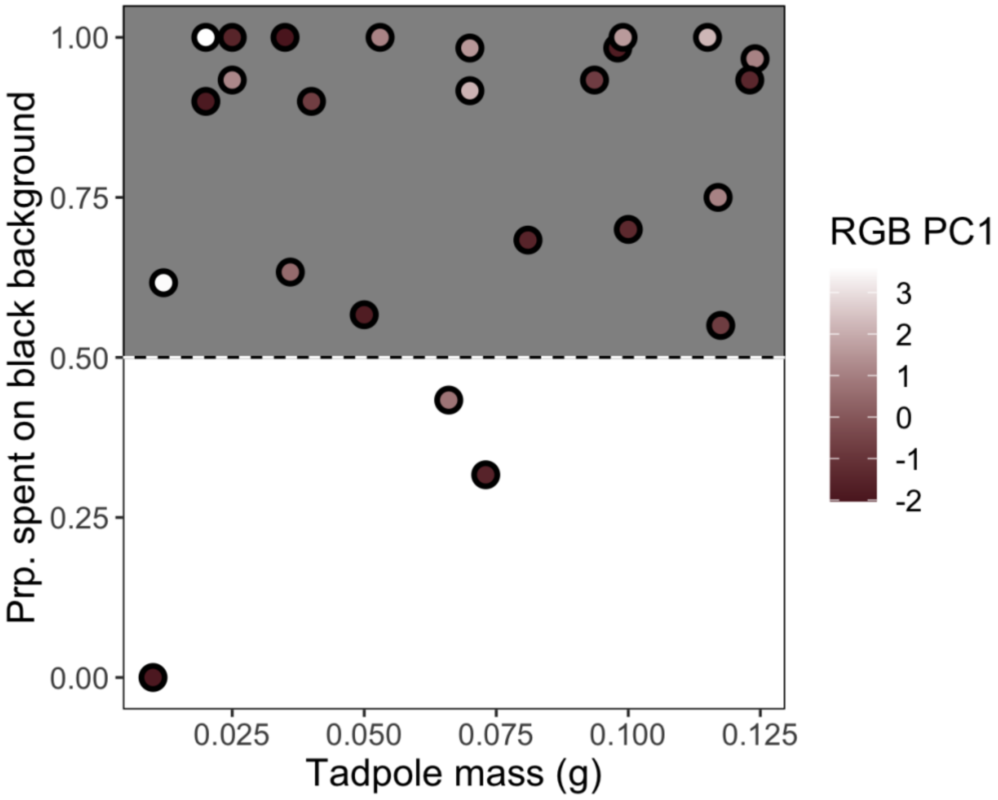

**Supplementary Figure 7.** Activity and focal tadpole mass correlation. In both species, we find a negative relationship between tadpole swimming and tadpole size (where size correlates to developmental stage in both species), where larger tadpoles tend to move less. Note the difference in scale on the x-axis between species.

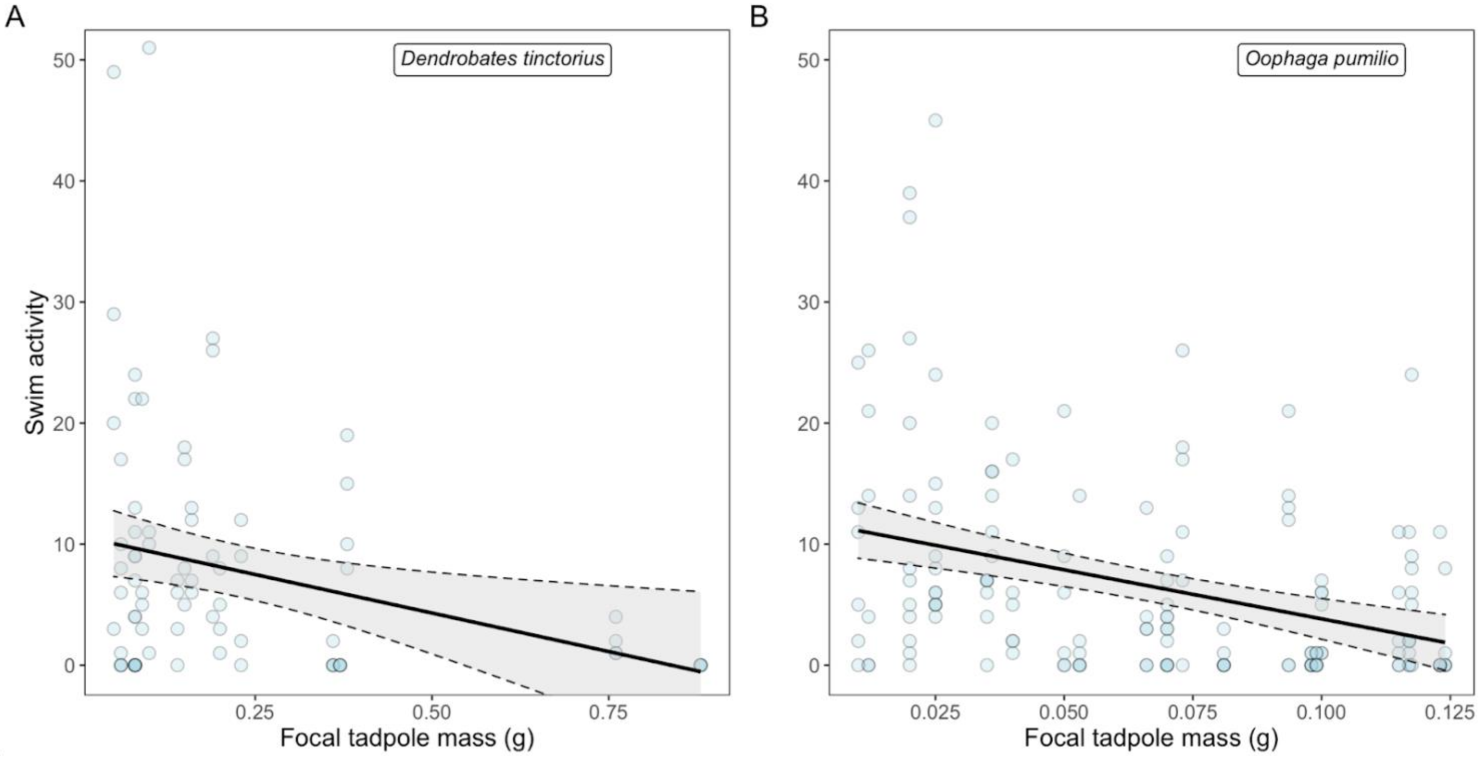


**Supplementary Figure 8.** Relationship between nursery conspecific counts and space use in *D. tinctorius*. Tadpoles from larval nurseries with more conspecifics spend less time in the arena centre. Interestingly, nursery conspecific count is highly correlated with the presence of predators (Pearson coefficient $r = 0.83$)

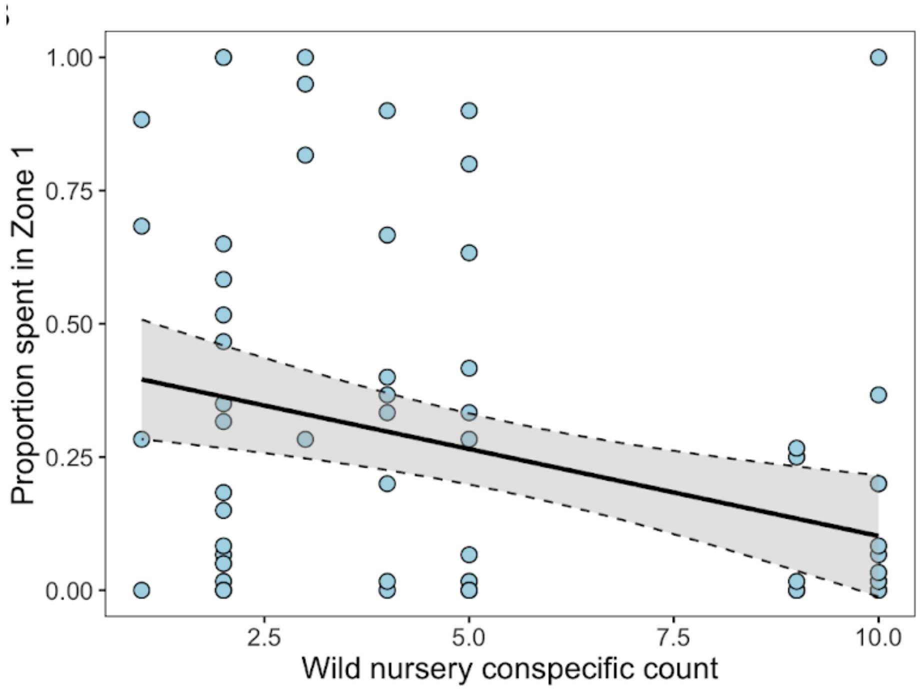
